# Expanding and enriching the LncRNA gene-disease landscape using the GeneCaRNA database

**DOI:** 10.1101/2024.03.11.584435

**Authors:** Shalini Aggarwal, Chana Rosenblum, Marshall Gould, Shahar Ziman, Ruth Barshir, Ofer Zelig, Yaron Guan-Golan, Tsippi Iny-Stein, Marilyn Safran, Shmuel Pietrokovski, Doron Lancet

**Author notes:** Correspondence: Shalini Aggarwal and Doron Lancet.

## Abstract

The GeneCaRNA human gene database is a member of the GeneCards Suite. It presents ∼280,000 human non-coding RNA genes, identified algorithmically from ∼690,000 RNAcentral’s transcripts. This expands by ∼tenfold the ncRNA gene count relative to other sources. GeneCaRNA thus contains ∼120,000 long non-coding RNAs (LncRNAs, >200 bases long), including ∼100,000 novel genes. The latter have sparse functional information, a vast terra incognita for future research. LncRNA genes are uniformly represented on all nuclear chromosomes, with 10 genes on mitochondrial DNA. Data obtained from MalaCards, another GeneCards Suite member, finds 1,547 genes associated with 1 to 50 diseases. ∼15% of the associations portray experimental evidence, with cancers tending to be multigenic. Preliminary text mining within GeneCaRNA discovers interactions of LncRNA transcripts with target gene products, with 25% being ncRNAs and 75% proteins. GeneCaRNA has a biological pathways section, which at present shows 131 pathways for 38 LncRNA genes, a basis for future expansion. Finally, our GeneHancer database provides regulatory elements for ∼110,000 LncRNA genes, offering pointers for co-regulated genes and genetic linkages from enhancers to diseases. We anticipate that the broad vista provided by GeneCaRNA will serve as an essential guide for further LncRNA research in disease decipherment.

## 1. Introduction

As our understanding of the human genome deepens, the spotlight on non-coding RNAs (ncRNAs) has intensified, revealing a diverse landscape of non-coding transcripts. Currently, the major sources (HGNC [1], NCBI Gene [2], Ensembl [3] and RNAcentral [4]), portray ∼38,000 well-documented ncRNA genes. In our earlier paper [5], we significantly augmented the count of identified ncRNA genes to 220,000, obtained based on algorithms that analyzed all transcripts in RNAcentral [4]. Our ncRNA gene category consists of 24 classes, with the highest counts belonging to LncRNAs, piRNAs and miRNAs.

GeneCaRNA’s *raison d’etre* within the GeneCards Suite is to provide a comprehensive gene-centric knowledgebase for human ncRNA genes and their annotations. The present version 5.19 GeneCaRNA exhibits ∼280,000 human non-coding RNA genes, stemming from ∼690,000 RNAcentral transcripts. A key advancement in GeneCaRNA is the addition of a very large number of potentially functional ncRNA genes to a gene da-tabase that enables the use of a broad range of other types of genomic and biological data and offers many bioinformatic analysis tools.

GeneCaRNA is a multi-source database, offering a promising avenue to enrich annotation information and functions on a single platform. GeneCaRNA effectively addresses the challenge of disparate gene names across sources by leveraging aliases, facilitating accurate gene mapping, and providing comprehensive gene details. GeneCaRNA is frequently updated to address novel knowledge and automatically add functional annotation and disease links which also address the novel Transcripts Inferred GeneCaRNA genes (TRIGGs)[5].

LncRNA genes constitute the largest class within the ncRNA gene category, ∼46% of the total. This gene class is defined for transcripts longer than 200 nucleotides, which have scant or no protein-coding characteristics [6,7]. The large size of the LncRNA class led to further definition of subclasses, based on criteria such as transcriptional machinery [8], structure [9], expression regulation [10], location with respect to protein coding genes [11,12], and binding targets [10], as well as transcriptional and post-transcriptional regulation [12].

LncRNA genes and their products are back-end regulators of the protein coding genes. These genes are majorly known to contribute to cancer pathobiology[13] and also to some hereditary diseases[14]. The mechanism of action is poorly understood, and their low transcription level makes them a covert disease driver[15]. Their role in non-cancerous diseases is yet to be discovered. We make some headway in this direction using GeneCaRNA and MalaCards by shedding light on the potential relations between LncRNAs and non-cancerous diseases.

This paper describes thorough analyses of the LncRNA class of human genes and their role in deciphering pathogenesis based on the power of the GeneCards Suite. GeneCaRNA provides textual analyses on annotations of each LncRNA GeneCard in the database and reveals information for items such as revised sub-classification, interactions between genes products (protein and transcripts), gene to disease relationships, and relations of genes to biological pathways. GeneCaRNA is poised to become a highly useful platform for accessing complete profiles of LncRNAs.

## 2. Methods

### 2.1. Data extraction and text mining

Data extraction from the GeneCaRNA database (GC V5.19, based on RNAcentral V23) employed Structured Query Language (SQL) or our in-house tool GeneAlaCart [16]. For routine analysis, we generated an interim LncRNA table containing the gene symbol, gene description, mined data sources, GeneCards Inferred Functionality Scores (GIFtS) [17], and chromosomal location of each gene (Table S1).

The extracted table contained 120,982 LncRNAs genes, whose categorization was based on transcript categorization in our sources. Some genes are assigned symbols based on the information in our major sources, and if absent there, are assigned our own GeneCards symbols as done before [16]. Most genes have additional (non-symbol) alias names featured in various external sources, which all are included in GeneCaRNA. GeneCaRNA was also used to extract all of the aliases and protein existence (PE) [18], using GeneAlaCart.

### 2.2. LncRNA Subclass definition

We extracted keywords from the Aliases section of GeneCaRNA, which contains multi-word gene descriptions. In the first stage, a list of the 10 most frequent words appearing in the entire collection of LncRNA genes was identified, using in-house python code (program code shown in S1 in **Supplementary file**). These terms were “Intronic”, “Intergenic”, “LINC”, “Overlapping”, “Divergent”, “Pseudogene”, “Antisense”, “Sense”, “Open Reading Frame”, and “ORF”. In a second stage, 5 candidate sub-classification terms were chosen, consulting other sub-classification approaches [3,11,19]. We performed sub-family term unification, e.g. “Overlapping”, “Pseudogene”, “Sense”, and “ORF” were clustered in our proposed system as “Protein Suspect”. Under the same sub-class, we also joined genes with the highest degree (1,2,3) of UniProt’s PE Levels [18], and the genes with lower levels (4,5) were also included provided that “protein” was a keyword appearing in our keyword search in GeneCaRNA’s Aliases section. Finally, each gene was assigned one or more sub-classes, based on the appearance of these keywords in the original gene/alias-specific text in the Aliases section of GeneCaRNA.

### 2.3. Gene-disease and Gene-Pathway associations

A list of diseases associated with each gene was obtained by an SQL query. Each gene-disease pairing was assigned an association score [16]. The score value depends on the level of manual curation of the information source, and on the significance assigned by the source itself to its different annotation classes. In addition, an “elite” grade is assigned for associations with experimental evidence support, and an “inferred” grade for the rest, where evidence is derived only from text-mining. GeneCards policy is to portray inferred association to catalyze future experimentation.

For gene-pathways relationships, we included only LncRNA genes with at least one associated disease. These were extracted using GeneAlaCart applied to the Pathways section of GeneCaRNA. The latter is mined from WikiPathways, SIGNOR, Reactome, PharmGKB, and Sino Biological pathways. In GeneCaRNA, we also show amalgamations to our SuperPathways facility [20].

### 2.4. Gene product Interactions

The interaction targets for each LncRNA transcript (proteins, ncRNAs or DNA segments) were identified by the Multigene Search (MuSe) in-house text mining program. The query gene can be a symbol or any of the aliases. This allows one to receive target genes for a long list of submitted genes, (e.g.) by submitting “<gene name> binds”, or “<gene name> interacts with”. The output is texts from e.g. the GeneCaRNA Publications or Summaries sections, that include the text query. The output file of the MuSe is further curated using in-house python code (see S2 in **Supplementary file**).

## 3. Results

### 3.1. Data extraction and text mining

We extracted 120,982 LncRNA genes from GeneCaRNA, as portrayed in **Table S1**. Of these, 19,482 genes are available in the major sources, and 101,500 genes are unique GeneCaRNA TRIGGs. Both categories were explored for annotative information based on their GIFtS value, which represents the cumulative information imported from all the relevant sources [17]. We found that while 34% of the major source genes had GIFtS > 10, none of the TRIGGs had a GIFtS > 10 (**Figure. 1**). This indicates the fertile land for future research in the realm of TRIGGs, both for LncRNA genes and other ncRNA categories.

**Figure 1.**
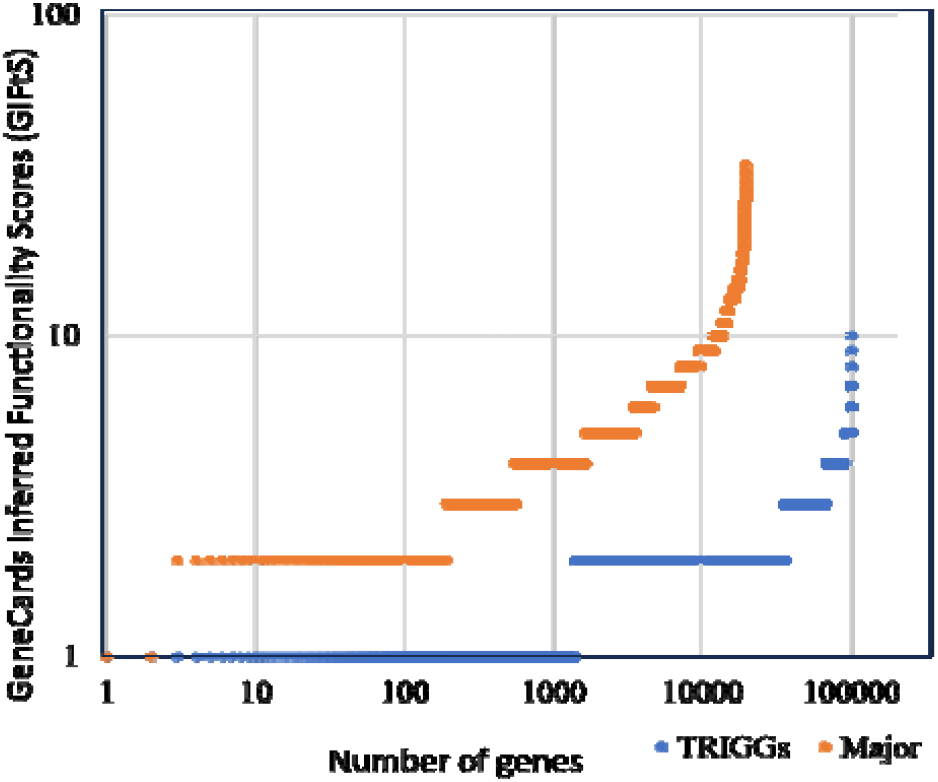
Rank graph for annotative information, separately illustrated for two groups: Genes from the major gene sources (NCBI, HGNC and ENSEMBL) (orange), and genes inferred from RNAcentral transcripts without any annotation from the major gene sources, TRIGGS (Blue).

### 3.2. LncRNA Subclass definition

We developed a pipeline for defining sub-classes for LncRNA genes (see Methods). [11,19]. We performed sub-family term unification (see Methods). As a result, five subclasses were created: “Antisense”, “LincRNA”, “Divergent”, “Protein Suspect”, and “Intronic” (**Table S2**). These sub-classes cover only 8.3% of the relevant genes; the rest were classified as “no subclass” (**Figure 2**).

**Figure 2.**
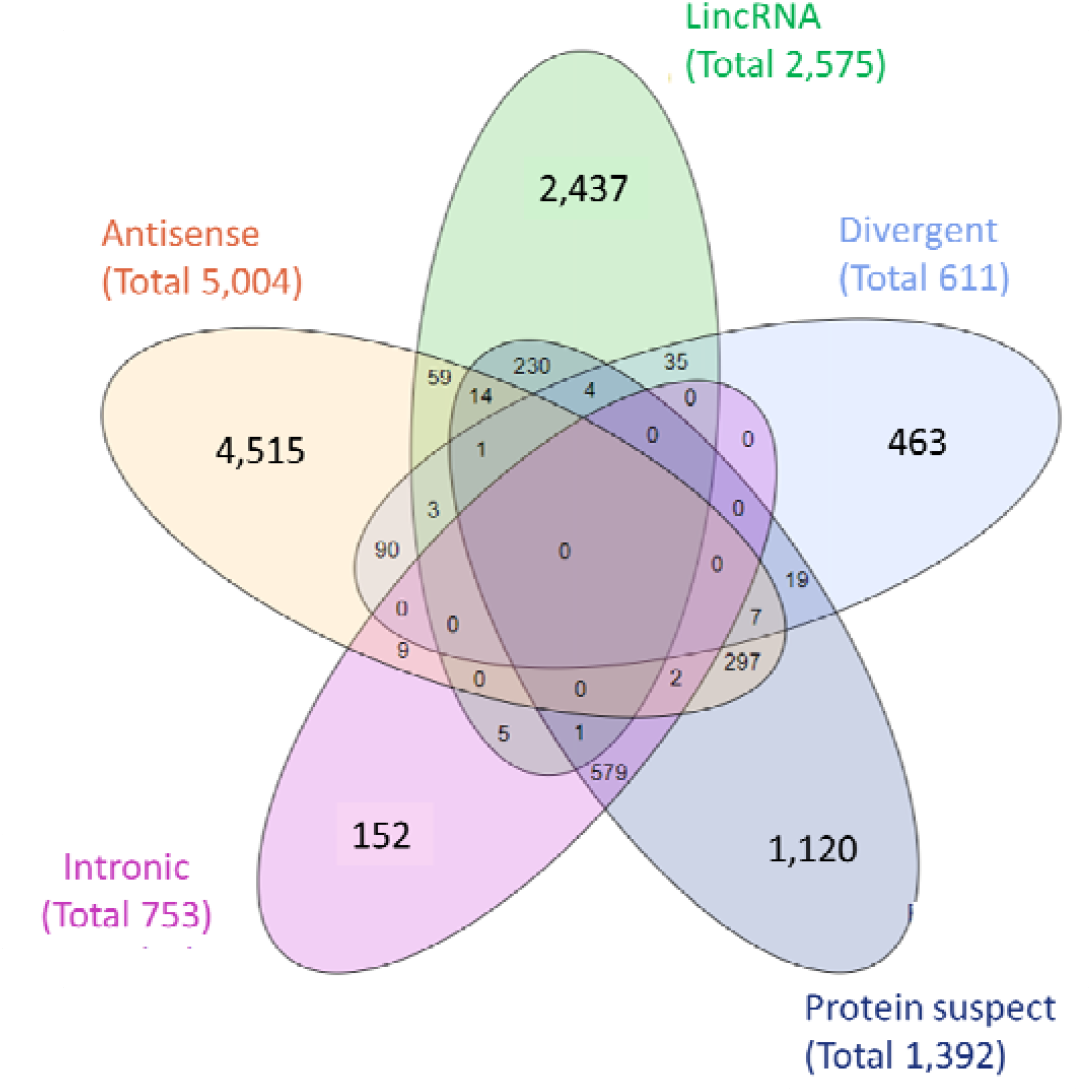
A GeneCaRNA-based suggested subclassification of 10,042 LncRNA subclass-definable genes (8.3% of the total). LincRNA: LncRNAs that are transcribed from the DNA stretch between the two protein coding genes. Divergent: LncRNAs that are transcribed by a promoter shared with a protein coding gene. Protein suspect: LncRNAs that have the potential to encode a peptide/protein. Intronic: LncRNAs transcribed purely from the intron(s) of a coding gene. Antisense: LncRNAs that are transcribed in antisense to a protein coding DNA strand.

### 3.3. Gene-disease and Gene-Pathway associations

We systematically explored GeneCaRNA for relationships of LncRNA genes to diseases, as portrayed in MalaCards, the disease database of the GeneCards suite [21]. A total of 5,100 gene-disease associations were found, spanning 1,554 LncRNA genes and 2,019 diseases. We identified 715 elite associations, involving 618 contributing diseases, and 340 contributing genes (**Table S3**). Among the 1,554 genes, 793 were associated with only 1 disease, and the rest had 2-111 associated diseases (**Figure 3a**). The top five disease-contributing genes are H19 (111 diseases), MALAT1 (87 diseases), MEG3 (76 diseases), HOTAIR (69 diseases), and CDKN2B-AS1 (59 diseases).

**Figure 3.**
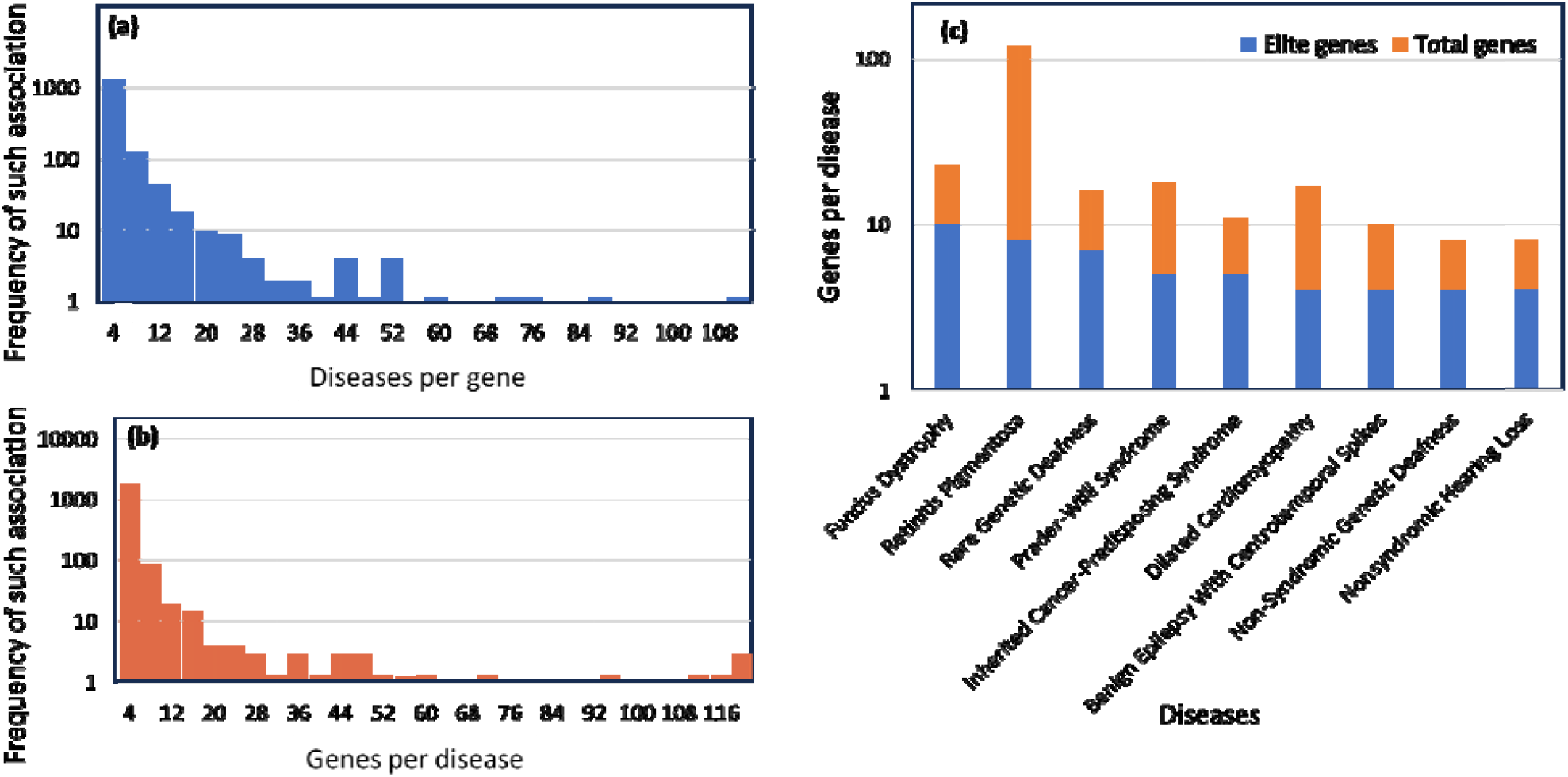
Gene-disease associations: Detailed scrutiny of the gene-disease associations among the 1,554 LncRNA genes and 2,019 diseases. **(a)** The count of diseases per gene values; **(b)** The count of gene per disease values; **(c)** The number of Genes per disease for a sample of 9 diseases with 4 or more elite associations.

Similarly, out of 2,019 diseases, 1,395 diseases associate with only 1 gene (**Figure 3b**). The top five diseases with the most genes are: Gastric cancer (120 genes), Retinitis pigmentosa (118 genes), Hepatocellular carcinoma (117 genes), Colorectal cancer (114 genes) and Lung cancer (111 genes). 1401 diseases with 3630 associations were inferred out of which 1047 diseases were with non-cancerous such as autism spectrum disorder (22 associations). Only 9 diseases were found to have 4 or more elite disease-gene associations; featured in **Figure 3c**. Around 50% of these genes are associated with diseases in many-to-many relationships, and the connectivity of selected cases of gene to disease associations are shown in **Figure 4**.

**Figure 4.**
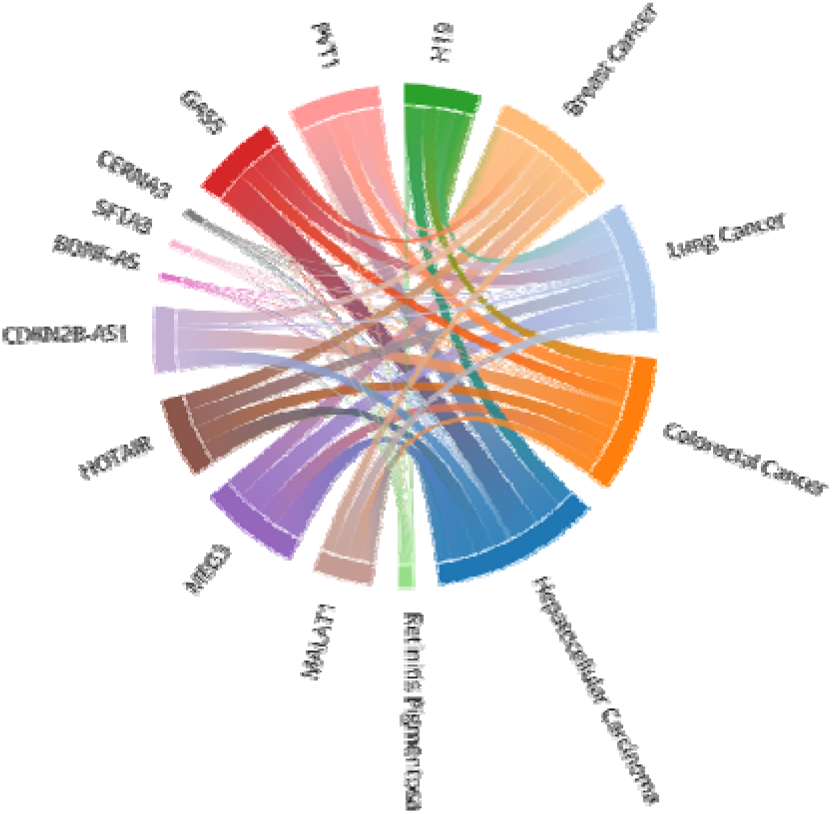
A map of genes to diseases associations. Chord diagram of the network of the top 10 most-recurring genes, and the top 5 most-recurring diseases. Each chord represents the gene/disease association, where the width of the line corelates with the association score. The association score follows the MalaCards gene to disease scoring system, where score values depend on the level of manual curation of the information source, and on the significance assigned by the source itself to its different annotation classes [21].

Involvement of multiple genes in a disease may indicate a gene-gene network, i.e. a biological pathway. Hence, all LncRNA genes associated with one or more disease were analyzed with respect to their pathway affiliation, as portrayed in the Pathways section of GeneCaRNA, which is based on the PathCards database in the GeneCards suite [20]. 131 pathways were found to be associated with 38 LncRNA genes (**Figures 5a** and **b and Table S4**). In an example, the role of four LncRNAs genes in a pathway with 28 genes is shown (**Figure 5c**).

**Figure 5.**
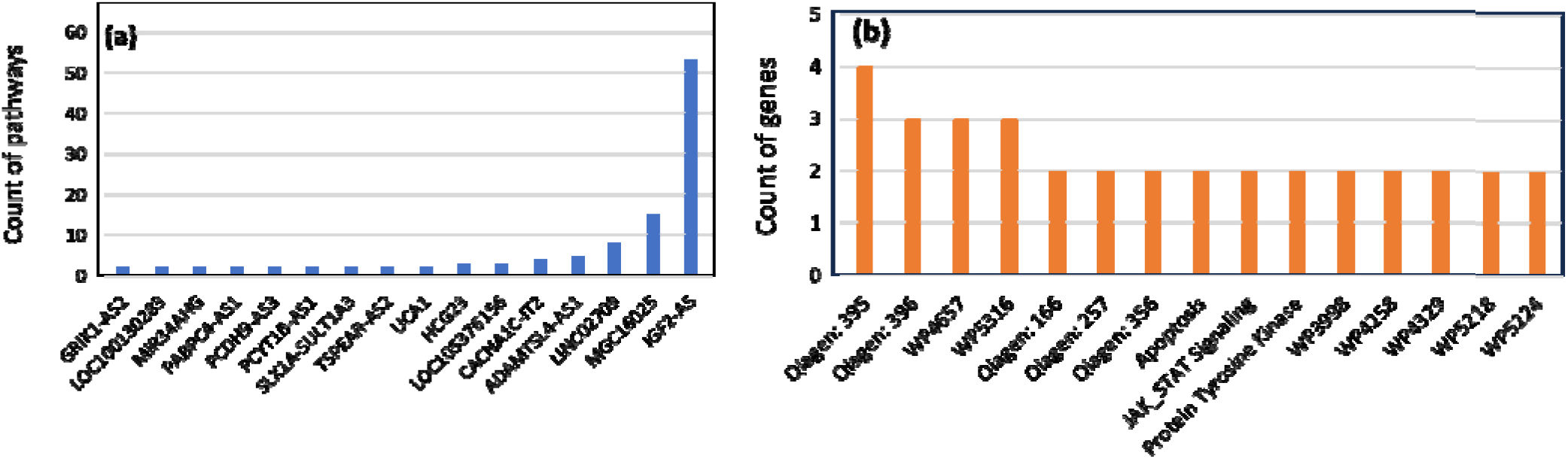
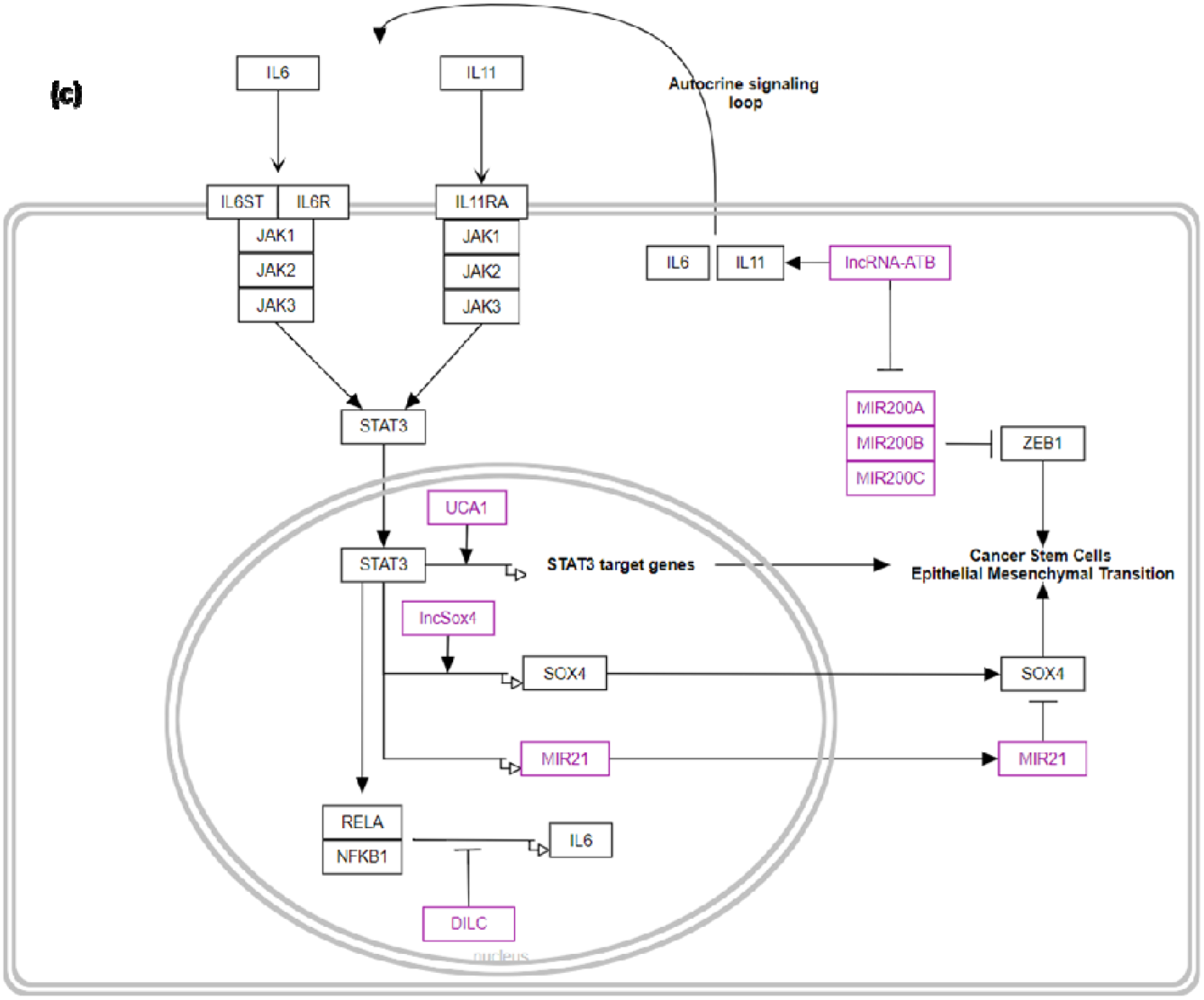
LncRNA-related pathways. **(a)** The counts of pathways (≥ 2) mapped to LncRNA genes; **(b)** The count of genes per pathway. 96 pathways include 1 gene (not shown) and at maximum a pathway had 4 genes; **(c)** A map of the “STAT3 signaling in hepatocellular carcinoma” obtained from WikiPathways (ID: WP4337). The LncRNA genes involved represent the cruciality of LncRNA genes in regulating protein coding genes, joining five MirRNA genes in the ncRNA category.

### 3.4. Gene product Interactions

Our multiple gene text analysis tool (MuSe) was applied to obtain a partial image of interactions of LncRNA transcripts with transcripts and encoded proteins of other genes. This was done with 3,000 adequately annotated LncRNA genes using 5 interaction terms. This facilitated the identification of 199 interacting partners mapped to 118 genes. These partners included 28 ncRNA genes and 171 protein/protein-coding genes. A sample of these interactions is shown in more detail in **Table 1**, showing the broad range of ncRNA interactions.

**Table 1.**
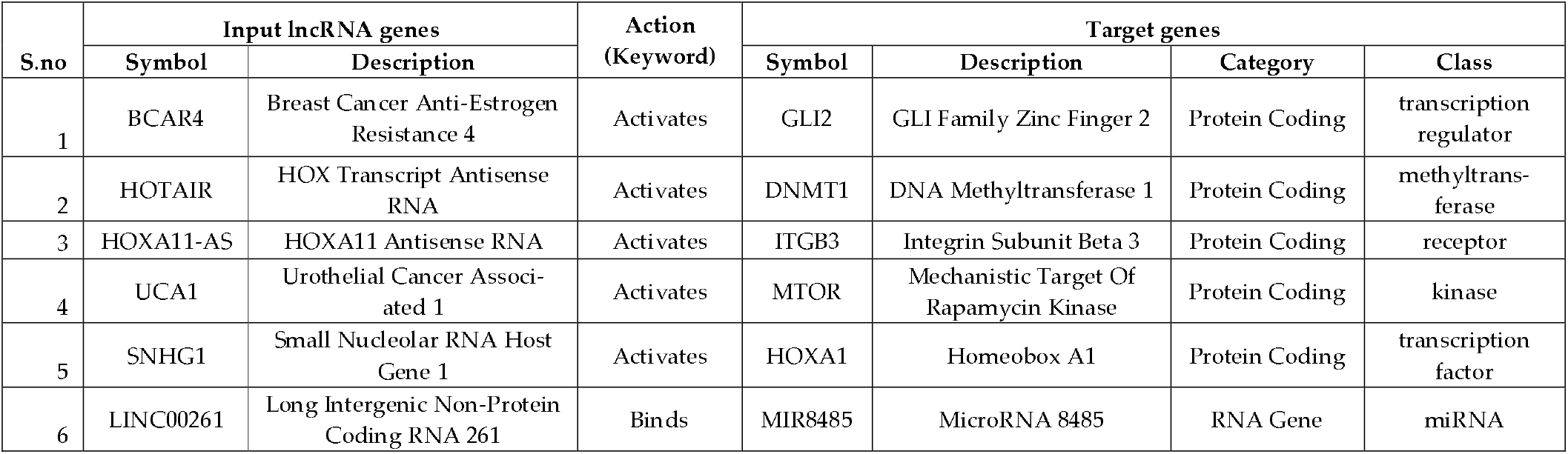
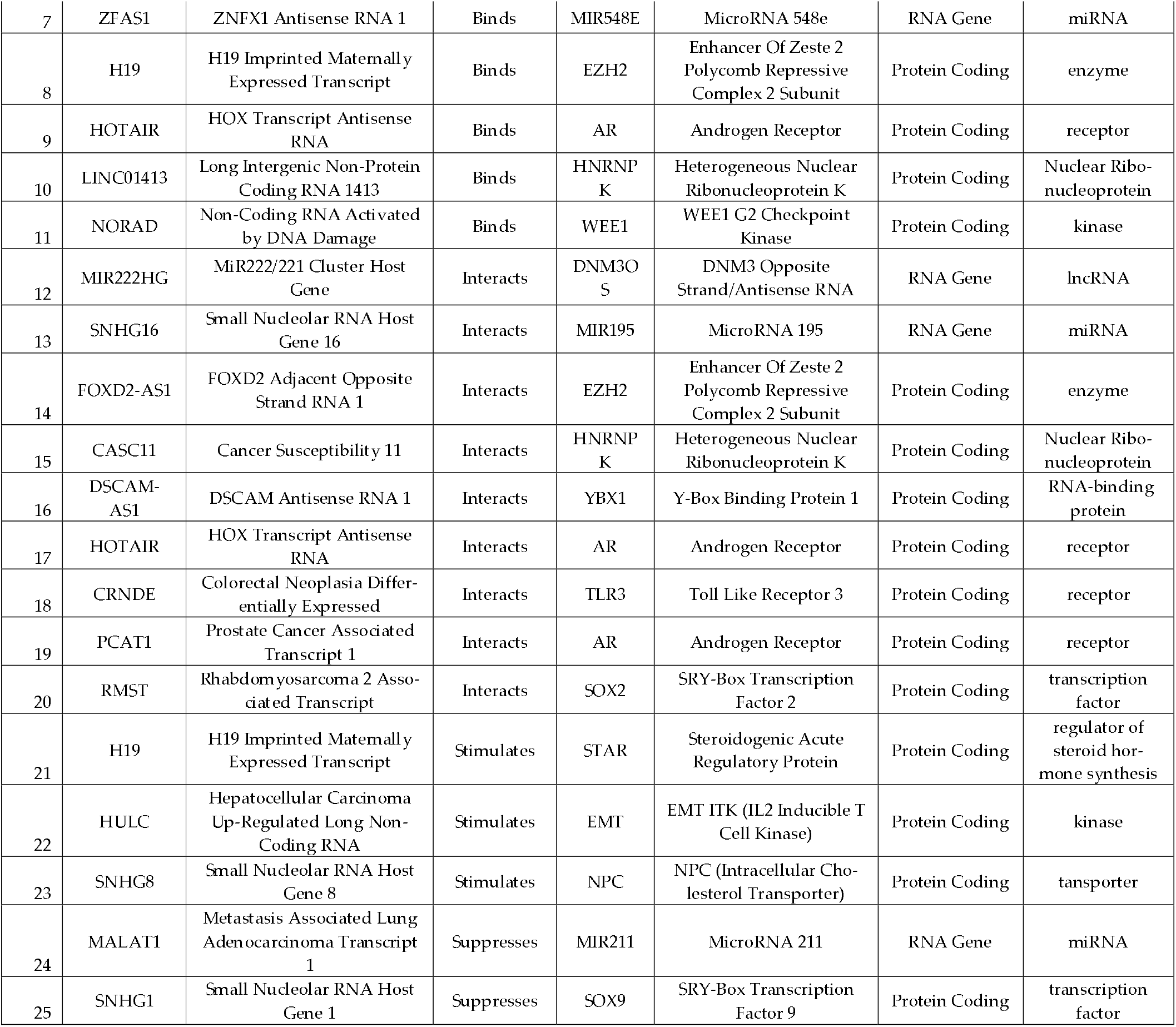
List of LncRNA genes and their associated target gene products.

Another method for finding functional relations is offered by GeneHancer, yet another facility of the GeneCards Suite [22]. This resource portrays regulatory elements (enhancers and promoters) for every gene in GeneCards, including all ncRNA genes. GeneHancer also provides a list of genes sharing enhancer(s) with the gene in question. It also exhibits potential phenotypes and diseases based on genome-wide association studies (GWAS) as shown in **Figure S1**. We extracted enhancer information for each LncRNA gene and found ∼108,000 genes having at least one enhancer mapped to it due to gene/enhancer proximity (**Figure S2**). 19,105 of the genes were contributed via major gene sources, with ∼11% having one-to-one relationship, and 88,974 were TRIGGs, with ∼17% of them in one-to-one relationship. Importantly, the very large group of novel TRIGG gene definitions often have no annotations except those inferred via GeneHancer, opening a way for a massive body of functional leading information.

**Figure 6.**
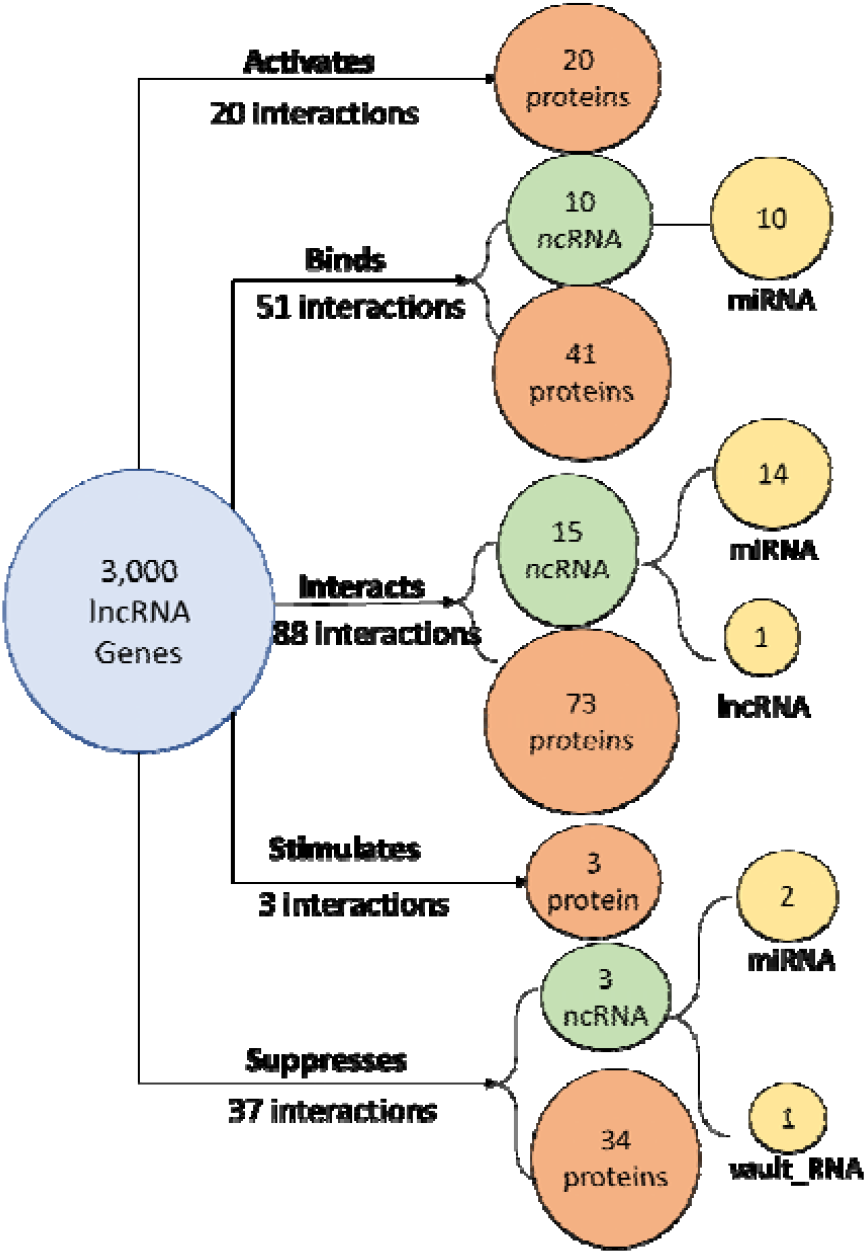
A diagram showing a sample of 199 target gene products showing one-to-one interactions with 118 LncRNA transcripts. The LncRNAs targets include 171 proteins, and a range of ncRNAs spanning three classes.

## 4. Discussion

GeneCaRNA was established 3 years ago to provide a compendium of all definable genes stemming from a full list of human transcripts. Its annotation system benefits from GeneCaRNA being part of the GeneCards Suite, which incorporates data-mined from 194 sources [16]. Our database succeeds in portraying as many as ∼280,000 ncRNA genes, ∼7-times more than several other resources. This significant improvement was attained by the TRIGG algorithm [16]. A great majority of the transcripts leading to the novel ∼240,000 genes are weakly annotated, but the richness of their hosting GeneCards Suite often provides new information for such genes. In this paper, we use this power to scrutinize the largest class of ncRNA genes, namely LncRNAs, a functionally diverse group of genes, which currently embodies an avalanche of new research. GeneCaRNA, encompassing ∼120,000 LncRNA genes, 43% of all ncRNA genes, provides substantial enrich-ment compared to other sources. Our GeneCaRNA-based enhancing pipeline for data extraction and analyses furnishes a fruitful approach to reporting and augment the relevant information published.

The immense size of the LncRNA class calls for sub-classification. The HUGO gene nomenclature committee (HGNC) defines six subclasses based on 6,000 LncRNAs [1]. On the other hand, GENCODE define nine sub-classes, based on 18,000 genes [11]. When analyzing all 120,000 GenCaRNA’s genes, we ascribed five subclasses to ∼10,000 genes, of which ∼600 were TRIGGS. We are now in the process of integrating GENCODE’s subclass assignments for 8,000 genes not yet covered by GeneCaRNA. All our five subclasses appear in other subclassification systems, whereby our sub-class “Protein suspect”, a seeming contradiction to the definition of LncRNA [23], is a merger of “3’ overlapping” and “Sense overlapping” in GENCODE. We elected not to include HGNC’s “contain microRNA or snoRNA” and GENCODE’s “macro lncRNA”, “non_coding” and “processed_transcript” LncRNA.

The study of LncRNAs is at the forefront of RNA biology due to the rise in reports claiming the role of LncRNAs in human disorders [24]. A key strength of GeneCaRNA is its inclusion in a framework that has profound information about diseases via the inter-database relations with MalaCards [21]. In parallel, there are mechanisms to discover indirect relationships between LncRNA genes and diseases via GeneHancer (Figure S2). Regarding the distinction between elite and inferred gene-disease associations, while the former are based on solid experimental support, the latter often provide a hint that opens a gateway for more research that may improve our understanding of a disease.

A key capacity of GeneCaRNA is text analyses on gene annotations, revealing information on interactions between ncRNA transcripts and other genes products (protein and transcripts). Using this platform, we were able to portray a partial view of such interaction network (**Table S5**). Further database development will allow us to obtain a holistic picture. Some examples of currently missing data include published functional roles of LncRNAs in the context of inflammatory diseases and cancers [25,26]. This happens by binding the LncRNAs to miRNAs, thus blocking their regulation of mRNAs in translation of downstream genes. Broadening the LncRNA interactome in GeneCaRNA will strengthen its capacity to support future research in the field.

The GeneCards suite includes VarElect [16,27], which can connect DNA variants to diseases. Initially, it was developed for exome sequencing, linking variants in protein coding genes to diseases. The establishment of GeneCaRNA now enables VarElect to function in the realm of disease decipherment of variants in ncRNAs genes obtained from whole genome sequencing. The indirect mode of VarElect helps discover association of variants in ncRNA genes with a disease known to be caused by mutations in a protein coding gene. This is achieved based on GeneCards information pointing to functional relationships between this ncRNA and the protein gene. This is important for solving undeciphered disease cases. This is an essential instrument for expanding the knowledge on the mechanisms by which LncRNA gene mutations lead to disease. In an example, somatic copy number mutations in lncRNA genes are shown to be associated with cancer clinical prognosis [28], where, at present, little is known about the molecular pathways involved.

GeneCaRNA provides a comprehensive unified repository of LncRNA and their annotative information. Exploited the forte of the GeneCards Suite, it is slated to become a powerful go-to place for researchers to explore these genes. The proposed text analysis pipeline for revealing gene-gene interactions to be integrated into the GeneCards platform, will likely improve the understanding of the role of LncRNAs in biological processes. The realm of the TRIGGS and inferred gene-disease associations are crucial terra incognita for pathobiology research especially for poorly understood diseases. As knowledge of LncRNAs continues to evolve, the frequently updating GeneCaRNA will serve as an essential platform, enabling researchers to navigate the expansive landscape of non-coding RNA, including LncRNAs, their targets, and their associated regulatory routes for disease decipherment.

## Supplementary Materials

The following supporting information can be downloaded at: www.mdpi.com/xxx/s1, Figure S1: Exhibits GeneHancer table; Figure S2: Rank graph to represent the distribution of number of enhancers per lncRNA gene; Table S1: List of all LncRNA genes in GeneCaRNA; Table S2: List of genes mapped to five sub-classes; Table S3: List of elite gene-disease associations; Table S4: List of gene-to-pathway associations; Table S5: List of gene-to-gene interactions for five selected keywords.

## Author Contributions

Conceptualization, D.L. and S.A.; methodology, D.L., S.A. and M.G.; software, S.A, C.R., M.G., O.Z., and Y.G.; validation, T.S. and S.A.; formal analysis, S.A.; investigation, S.A.; resources, C.R., O.Z., Y.G.; data curation, S.A.; writing—original draft preparation, S.A.; writing—review and editing, M.S., S.P., M.G., and D.L.; visualization, S.A., and S.Z.; supervision, D.L.; funding acquisition, D.L. All authors have read and agreed to the published version of the manuscript.

## Institutional Review Board Statement

Not applicable.

## Informed Consent Statement

Not applicable.

## Data Availability Statement

Use of GeneCards Suite websites is free for academic non-profit institutions. The suite’s extensive knowledgebase is available for research purposes via an academic collaboration agreement (see https://www.genecards.org/Guide/Datasets), Other users must acquire a <u>commercial license from LifeMap Sciences</u>.

## Acknowledgments

None

## Appendix A

- LncRNA: Long non-coding RNA
- ncRNA: non-coding RNA
- HGNC: HUGO Gene Nomenclature Committee
- NCBI: National Center for Biotechnology Information
- TRIGGs: Transcripts Inferred GeneCaRNA genes.
- SQL: Structured Query Language
- GIFtS: GeneCards Inferred Functionality Scores
- LincRNA: Long intergenic non-coding RNA
- MuSe: Multigene Search
- ORF: Open Reading Frame
- PE: Protein Evidence

## References

1. Seal, R.L.; Chen, L.-L.; Griffiths-Jones, S.; Lowe, T.M.; Mathews, M.B.; O’Reilly, D.; Pierce, A.J.; Stadler, P.F.; Ulitsky, I.; Wolin, S.L.; et al. A Guide to Naming Human Non-Coding RNA Genes. EMBO J. 2020, 39, e103777, doi:10.15252/embj.2019103777.

2. Brown, G.R.; Hem, V.; Katz, K.S.; Ovetsky, M.; Wallin, C.; Ermolaeva, O.; Tolstoy, I.; Tatusova, T.; Pruitt, K.D.; Maglott, D.R.; et al. Gene: A Gene-Centered Information Resource at NCBI. Nucleic Acids Res. 2015, 43, D36–42, doi:10.1093/nar/gku1055.

3. Harrison, P.W.; Amode, M.R.; Austine-Orimoloye, O.; Azov, A.G.; Barba, M.; Barnes, I.; Becker, A.; Bennett, R.; Berry, A.; Bhai, J.; et al. Ensembl 2024. Nucleic Acids Res. 2024, 52, D891–D899, doi:10.1093/nar/gkad1049.

4. The RNAcentral Consortium; Sweeney, B.A.; Petrov, A.I.; Burkov, B.; Finn, R.D.; Bateman, A.; Szymanski, M.; Karlowski, W.M.; Gorodkin, J.; Seemann, S.E.; et al. RNAcentral: A Hub of Information for Non-Coding RNA Sequences. Nucleic Acids Res. 2019, 47, D221–D229, doi:10.1093/nar/gky1034.

5. Barshir, R.; Fishilevich, S.; Iny-Stein, T.; Zelig, O.; Mazor, Y.; Guan-Golan, Y.; Safran, M.; Lancet, D. GeneCaRNA: A Comprehensive Gene-Centric Database of Human Non-Coding RNAs in the GeneCards Suite. J. Mol. Biol. 2021, 433, 166913, doi:10.1016/j.jmb.2021.166913.

6. Kapranov, P.; Cheng, J.; Dike, S.; Nix, D.A.; Duttagupta, R.; Willingham, A.T.; Stadler, P.F.; Hertel, J.; Hackermüller, J.; Hofacker, I.L.; et al. RNA Maps Reveal New RNA Classes and a Possible Function for Pervasive Transcription. Science 2007, 316, 1484–1488, doi:10.1126/science.1138341.

7. Wang, J.; Zhu, S.; Meng, N.; He, Y.; Lu, R.; Yan, G.-R. ncRNA-Encoded Peptides or Proteins and Cancer. Mol. Ther. 2019, 27, 1718, doi:10.1016/j.ymthe.2019.09.001.

8. Mattick, J.S.; Amaral, P.P.; Carninci, P.; Carpenter, S.; Chang, H.Y.; Chen, L.-L.; Chen, R.; Dean, C.; Dinger, M.E.; Fitzgerald, K.A.; et al. Long Non-Coding RNAs: Definitions, Functions, Challenges and Recommendations. Nat. Rev. Mol. Cell Biol. 2023, 24, 430–447, doi:10.1038/s41580-022-00566-8.

9. Aliperti, V.; Skonieczna, J.; Cerase, A. Long Non-Coding RNA (lncRNA) Roles in Cell Biology, Neurodevelopment and Neurological Disorders. Non-Coding RNA 2021, 7, 36, doi:10.3390/ncrna7020036.

10. Ma, L.; Bajic, V.B.; Zhang, Z. On the Classification of Long Non-Coding RNAs. RNA Biol. 2013, 10, 924–933, doi:10.4161/rna.24604.

11. Frankish, A.; Diekhans, M.; Jungreis, I.; Lagarde, J.; Loveland, J.E.; Mudge, J.M.; Sisu, C.; Wright, J.C.; Armstrong, J.; Barnes, I.; et al. GENCODE 2021. Nucleic Acids Res. 2021, 49, D916–D923, doi:10.1093/nar/gkaa1087.

12. Sun, Q.; Hao, Q.; Prasanth, K.V. Nuclear Long Noncoding RNAs: Key Regulators of Gene Expression. Trends Genet. TIG 2018, 34, 142–157, doi:10.1016/j.tig.2017.11.005.

13. Lekka, E.; Hall, J. Noncoding RNAs in Disease. Febs Lett. 2018, 592, 2884–2900, doi:10.1002/1873-3468.13182.

14. Sparber, P.; Filatova, A.; Khantemirova, M.; Skoblov, M. The Role of Long Non-Coding RNAs in the Pathogenesis of Hereditary Diseases. BMC Med. Genomics 2019, 12, 42, doi:10.1186/s12920-019-0487-6.

15. Zhang, X.; Wang, W.; Zhu, W.; Dong, J.; Cheng, Y.; Yin, Z.; Shen, F. Mechanisms and Functions of Long Non-Coding RNAs at Multiple Regulatory Levels. Int. J. Mol. Sci. 2019, 20, 5573, doi:10.3390/ijms20225573.

16. Safran, M.; Rosen, N.; Twik, M.; BarShir, R.; Stein, T.I.; Dahary, D.; Fishilevich, S.; Lancet, D. The GeneCards Suite. In Practical Guide to Life Science Databases; Abugessaisa, I., Kasukawa, T., Eds.; Springer Nature: Singapore, 2021; pp. 27–56 ISBN 9789811658129.

17. Harel, A.; Inger, A.; Stelzer, G.; Strichman-Almashanu, L.; Dalah, I.; Safran, M.; Lancet, D. GIFtS: Annotation Landscape Analysis with GeneCards. BMC Bioinformatics 2009, 10, 348, doi:10.1186/1471-2105-10-348.

18. Lane, L.; Bairoch, A.; Beavis, R.C.; Deutsch, E.W.; Gaudet, P.; Lundberg, E.; Omenn, G.S. Metrics for the Human Proteome Project 2013-2014 and Strategies for Finding Missing Proteins. J. Proteome Res. 2014, 13, 15–20, doi:10.1021/pr401144x.

19. Seal, R.L.; Tweedie, S.; Bruford, E.A. A Standardised Nomenclature for Long Non-Coding RNAs. IUBMB Life 2023, 75, 380–389, doi:10.1002/iub.2663.

20. Belinky, F.; Nativ, N.; Stelzer, G.; Zimmerman, S.; Iny Stein, T.; Safran, M.; Lancet, D. PathCards: Multi-Source Consolidation of Human Biological Pathways. Database J. Biol. Databases Curation 2015, 2015, bav006, doi:10.1093/database/bav006.

21. Rappaport, N.; Twik, M.; Plaschkes, I.; Nudel, R.; Iny Stein, T.; Levitt, J.; Gershoni, M.; Morrey, C.P.; Safran, M.; Lancet, D. MalaCards: An Amalgamated Human Disease Compendium with Diverse Clinical and Genetic Annotation and Structured Search. Nucleic Acids Res. 2017, 45, D877–D887, doi:10.1093/nar/gkw1012.

22. Fishilevich, S.; Nudel, R.; Rappaport, N.; Hadar, R.; Plaschkes, I.; Iny Stein, T.; Rosen, N.; Kohn, A.; Twik, M.; Safran, M.; et al. GeneHancer: Genome-Wide Integration of Enhancers and Target Genes in GeneCards. Database J. Biol. Databases Curation 2017, 2017, bax028, doi:10.1093/database/bax028.

23. Mattick, J.S.; Rinn, J.L. Discovery and Annotation of Long Noncoding RNAs. Nat. Struct. Mol. Biol. 2015, 22, 5–7, doi:10.1038/nsmb.2942.

24. Wapinski, O.; Chang, H.Y. Long Noncoding RNAs and Human Disease. Trends Cell Biol. 2011, 21, 354–361, doi:10.1016/j.tcb.2011.04.001.

25. Lin, Y.; Jin, L.; Tong, W.M.; Leung, Y.Y.; Gu, M.; Yang, Y. Identification and Integrated Analysis of Differentially Expressed Long Non-coding RNAs Associated with Periodontitis in Humans. J. Periodontal Res. 2021, 56, 679–689, doi:10.1111/jre.12864.

26. Salmena, L.; Poliseno, L.; Tay, Y.; Kats, L.; Pandolfi, P.P. A ceRNA Hypothesis: The Rosetta Stone of a Hidden RNA Language? Cell 2011, 146, 353–358, doi:10.1016/j.cell.2011.07.014.

27. Stelzer, G.; Plaschkes, I.; Oz-Levi, D.; Alkelai, A.; Olender, T.; Zimmerman, S.; Twik, M.; Belinky, F.; Fishilevich, S.; Nudel, R.; et al. VarElect: The Phenotype-Based Variation Prioritizer of the GeneCards Suite. BMC Genomics 2016, 17, 444, doi:10.1186/s12864-016-2722-2.

28. Du, Z.; Fei, T.; Verhaak, R.G.W.; Su, Z.; Zhang, Y.; Brown, M.; Chen, Y.; Liu, X.S. Integrative Genomic Analyses Reveal Clinically Relevant Long Noncoding RNAs in Human Cancer. Nat. Struct. Mol. Biol. 2013, 20, 908–913, doi:10.1038/nsmb.2591.

