## Supplementary material for "Expanding and enriching the LncRNA gene-disease landscape using the GeneCaRNA database": Supplemental_Fig_S1.pptx

### Slide 1
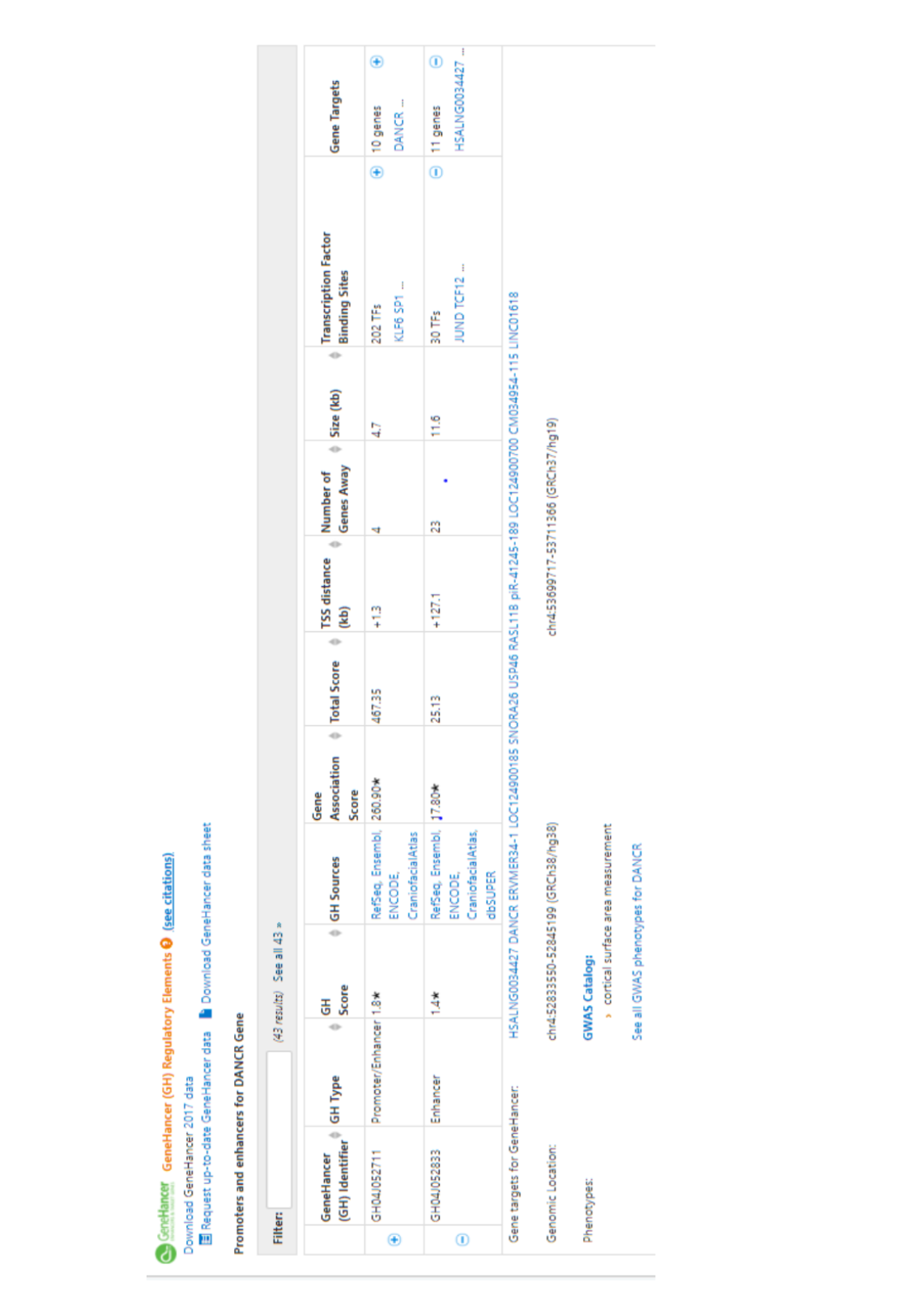

### Slide 2
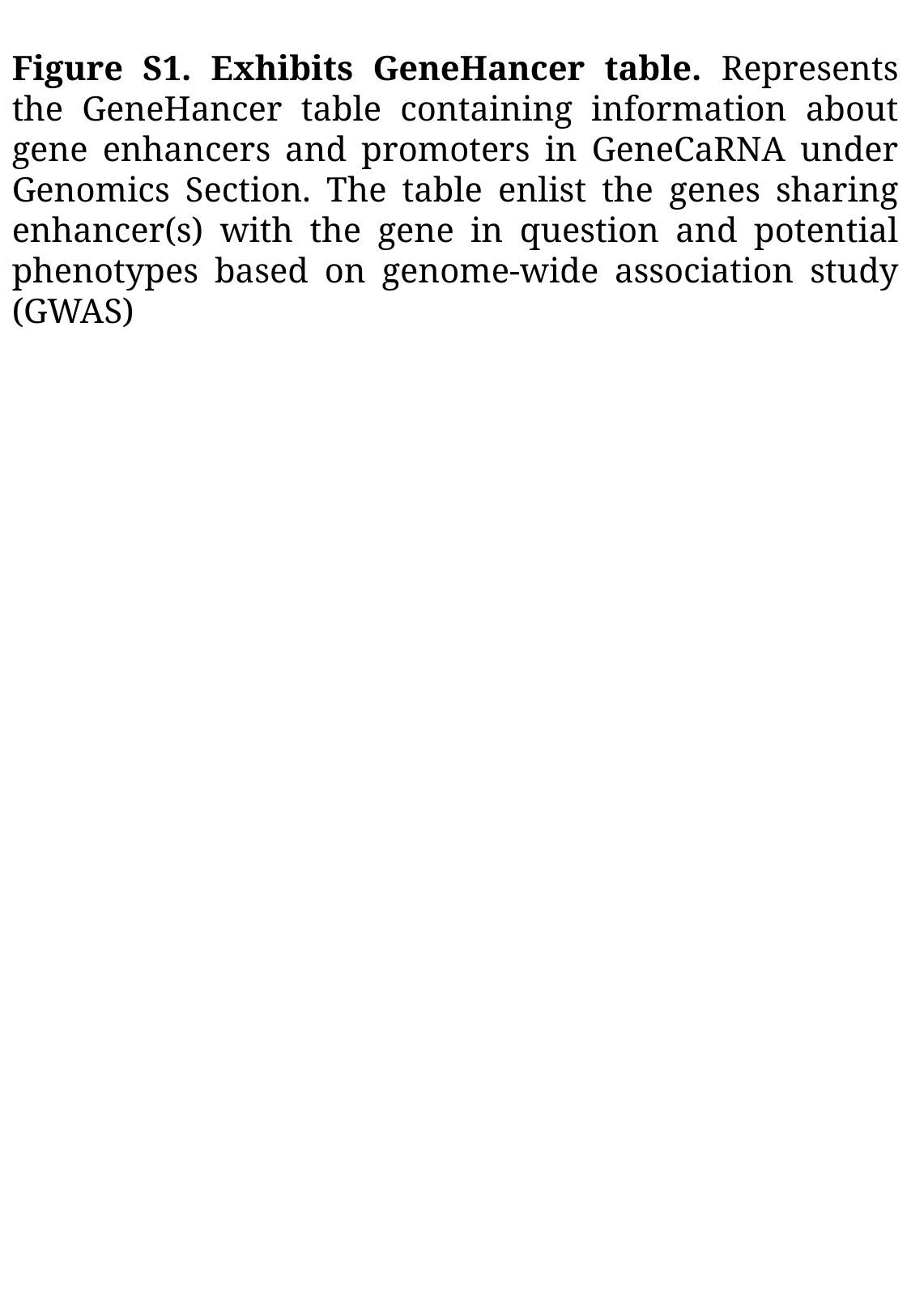

Figure S1. Exhibits GeneHancer table. Represents the GeneHancer table containing information about gene enhancers and promoters in GeneCaRNA under Genomics Section. The table enlist the genes sharing enhancer(s) with the gene in question and potential phenotypes based on genome-wide association study (GWAS)
