## Supplementary material for "Expanding and enriching the LncRNA gene-disease landscape using the GeneCaRNA database": Supplemental_Fig_S2.pptx

### Slide 1
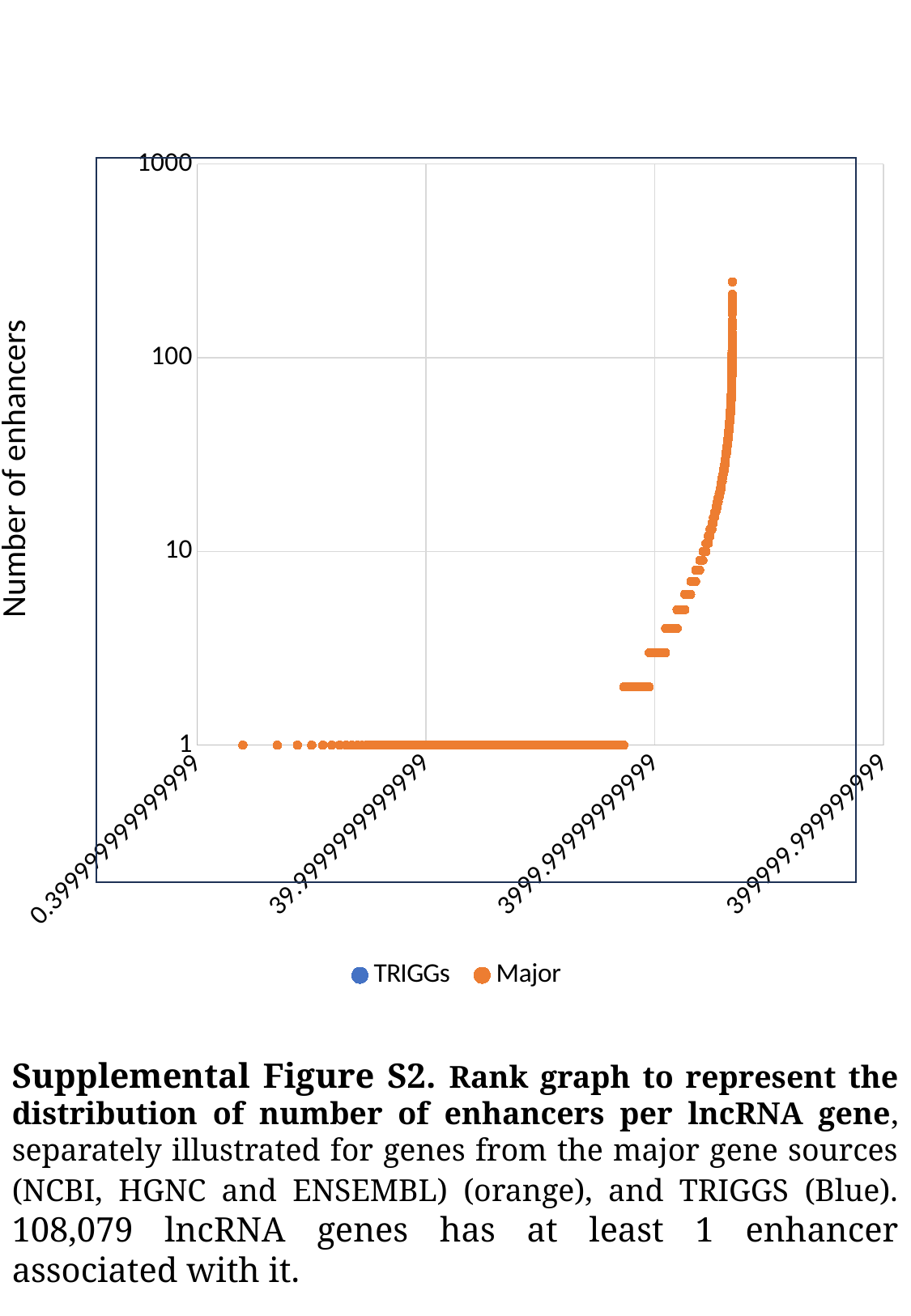

#### Chart
| Category | TRIGGs | Major |
|---|---|---|
Number of enhancers
Supplemental Figure S2. Rank graph to represent the distribution of number of enhancers per lncRNA gene, separately illustrated for genes from the major gene sources (NCBI, HGNC and ENSEMBL) (orange), and TRIGGS (Blue). 108,079 lncRNA genes has at least 1 enhancer associated with it.
