## Supplementary material for "Expanding and enriching the LncRNA gene-disease landscape using the GeneCaRNA database": Supplemental_Methods.docx

^3^ TAD Center for AI and Data Science, Tel Aviv University, Tel Aviv 6997801, Israel

**S.1.** **Code used for text analysis on Python 3.10.** This analysis leverages Python's standard libraries, notably the collections module for its Counter class, to efficiently tally word frequencies in text data. The code used for the two steps are mentioned below:

1. The frequency of each word within the aliases was determined, and they were then ranked in descending order of their frequency. The code used for this analysis was:

simple_keyword_counter = Counter()

for alias in unique_meaningful_aliases:

words = alias.lower().split()

simple_keyword_counter.update(words)

sorted_simple_keywords =

sorted(simple_keyword_counter.items(), key=lambda x: x[1], reverse=True)

2. Reordering of Aliases Based on Keyword Frequency:

Each alias was reordered based on the frequency of its constituent keywords. For instance, if an alias is "Carcinoma Transcript", and "Transcript" appears more frequently than "Carcinoma" in the list of aliases, the reordered alias would be "Transcript Carcinoma".

def reorder_alias(alias, keyword_freq_dict):

words = alias.split()

reordered_words =

sorted(words, key=lambda x: keyword_freq_dict.get(x.lower(), 0), reverse=True)

return ' '.join(reordered_words)

reordered_aliases =

[reorder_alias(alias, simple_keyword_counter) for alias in unique_meaningful_aliases]

sorted_reordered_aliases = sorted(reordered_aliases)

**S.2.** **Python script was used to generate CSV file from an HTML file** (Output of MuSe), showcasing LncRNA genes and their identified interactors from the GeneCards Suite.

**Summary of the logic flow:**

1. **MuSe output file processing:**
   - Identified and extracted the search query and paragraphs from the MuSe output file. Query genes were the selected 3000 LncRNA genes.
   - Empty search results were removed.
   - Unique Sentences Generation: For each query gene, mapped paragraphs were reduced to five words length starting from the five action keywords i.e. suppresses, interacts, activates, stimulates, and binds.
2. **Gene Information Extraction from GC Suite:**
   - Gene CSV generation: For each Gene symbol in the GC Suite aliases, category, class and description were extracted in CSV format.
4. **Gene Database Integration:**
   - Unique Sentences were screened for gene symbols and mapped against Gene CSV file for information about the gene symbol, category, and class.
5. **Data Cleanup and Filtering:**
   - The merged information was filtered and sorted to remove irrelevant information.
6. **Exporting Results:**
   - The final dataframe, containing relevant gene information and sentences, was exported to a CSV file **(Table S5)**.

Code can be accessed: <https://github.com/ShaliniAggy/MuSe-OP-analysis>
